# The role of coherence in bacterial communication

**DOI:** 10.1101/119503

**Authors:** Sarangam Majumdar, Sisir Roy

## Abstract

Bacteria within biofilms can coordinate their behavior through distinct from of communication mechanism^1^. The well-established cell - to - cell signaling process in bacteria is known as quorum sensing through chemical signaling molecules^2-5^. Recently, another cell- to - cell communication process based on ion channel mediated electrical signaling^6^ has also been observed. In this article, we propose a novel approach to explain the role of coherence and phase synchronization in the cell – to – cell bacterial communication. The observable long – range coherent electrical signaling is species independent and it is caused by membrane – potential - dependent modulation of tumbling frequency^7-9^. Moreover, noise can play a constructive role in enhancing the synchronization of chaotic bacterial communication systems and noise associated with the opening and closing the gate of ion channel induce small kinetic viscosity that make a wave-like pattern in concentration profile of quorum sensing.

---

The essence of mathematics and physics are mainly represented by its enormous capability to understand any complex natural phenomenon, from the evolution of life to the origin of universe. Every biochemical systems are composed of complex microscopic and macroscopic phenomena, which are very hard to explain, by a set of pure classical mathematical equations. Thus so far simple physical principals have remained unable in describing all the notions that came out through these biochemical machineries^10^. Quorum sensing mechanism is the best known charecteristic of the cell - to -cell communication in the bacterial world. Quorum sensing bacteria emit a singnaling molecules which is known as autoinducer (AI- 1 and AI-2)^2-4^. However, do not consider them to be small and simple, because they possess the generic term quorum sensing adopted to describe the cell communication process which coordinate gene expression, when the population has reached a high cell density. There are many situations when the bacterial population behave cooperatively and recognize self and non- self which can be highly advantageous, particularly in the contexts of sex, symbiosis, niche adaptation, production of secondary metabolites combined with the defense mechanisms of higher organisms and for facilitating population migration where the prevailing conditions in a specific environmental niche have become unfavorable^5^.

For several years, the study of bacterial ion channels has provided fundamental insights into the structural basis of such neuronal signalling^11^. It has been observed that bacteria process many important classes of ion channels, such as sodium channeles^12^, cholride channels^13^, calcium- gated potassium channels^14^ and ionotropic glutamate receptors^15^. But the significant role of these bacterial ion channels are still unclear. In 2015, it has been observed that ion channels conduct long- range electrical signals within the bacterial biofilm communities through spatially propagating waves of potassium^6^. Moreover, it has been shown that K^+^ ion channel mediated electrical signaling generated by *B. subtilis* biofilm can attract distant cells, thereby directing their mobility^14^.

Bacteria residing within biofilm communities can coordinate their behaviour through cell-to-cell signaling. Recently, another cell -to -cell signaling mechanism is discovered which is based on ion channel mediated electrical signaling. This electrical signaling has been shown to facilitate communication within a biofilm community^11^. Bacterial cells within *B. subtilis* biofilms can actively relay extracellular potassium signals, producing electrical waves that propagate through the biofilm and coordinate metabolic states, thereby increasing collective fitness^6-8^. Humphries et al.^9^ find that potassium ion channel mediated electrical signaling generated by *B. subtilis* bioflim can attract distant cells. Experimental and mathematical modeling indicates that extracellular potassium emitted from the biofilm alters the membrane potential of distant cells thereby directing their motility. Moreover, this electrically mediated attraction is generic in nature that enables crossspecies communication, as *P. aeruginosa*(quorum sensing bacteria) cells also become attracted to the electrical signal released by *B. subtilis* biofilm.

Adherent communities of *Bacillus subtilis* form biofilms and grow in interval of cycles once the colony reaches threshold size of population. These cycles arise when the cells present in the biofilms rundown of glutamate due to consumption of high amount of amino acid by peripheral cells. Glutamate starvation in the interior cells reduces the production of ammonium ions, which is required by the peripheral cells. As a result, the cell growth diminishes drastically^7^. Mathematical models on signaling explained to an extent the question raised on how following linked metabolic processes of cells within the biofilm community supports a distant communication. Cells within a biofilm community can thus not only coordinate their behaviour but also influence the behaviour of diverse bacteria at a distance through long-range electrical signaling^9^.

It has been noted in the experiment that oscillation in membrane potential were synchronized among even the most distant regions of the biofilm community^6^. It raises the important question whether active electrochemical signaling could be responsible for the long - range synchronization or not. Motivated from the recent experimental evidences of species independent attraction to the biofilms through electrical signaling, we can predict some important role of coherence and phase synchronization. We propose the important role of noise in producing the coherence and synchronization. This noise is due to the opening and closing of gate of potassium ion channel. As the cell density reaches to particular threshold density, quorum sensing occurs. The highly dense bacterial colony is associated with many potassium ion channels. So it gives rise threshold noise to produce synchronization and hence the coherence.

We are interested to investigate the interrelation in the quorum sensing (through chemical signaling) mechanism and the electrical communication in bacterial communities. The detection of such interactions enables deeper insight into the basic process and the functioning underlying these systems. The nonparametric coherence analysis developed in the framework of the linear stochastic systems enables a detection of interactions in transfer function systems. It is very sensitive in detecting interactions in several systems^16^. From the coherence value, it is not possible to conclude something about the underlying dynamics. We can consider the oscillation in membrane potential as nonlinear synchronizing oscillators (bacterial cell membrane) and phase synchronization analysis has been shown to be very efficient in detecting interactions between oscillators.

In order to detect phase synchronization between coupled oscillatory systems, a suitable definition of phase and amplitude of the real valued observed potassium ion channel mediated signal is necessary. Phase synchronization of coupled, chaotic oscillations occurs if the phase lock condition is satisfied^17^. For the phase jump induced by the presence of dynamics or observation noise is modified. Let us consider *Ψ* (*t*) to be an analytic electrical signal and *n, m* be the positive integers. In that situation, a sharp peak in the distribution of *Ψ*_*n, m*_ indicates phase-synchronized oscillators. Then, we can find the synchronized index 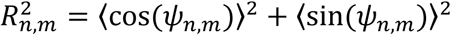. If *R*_*n,m*_ = 1 then we say that a constant phase difference in oscillators and *R*_*n, m*_ = 0 for uniformly distributed phase difference.

So, phase synchronization analysis is not specific in detecting the correct class of underling dynamics, as phase synchronization is based on coupled self-sustained oscillators. On the other hand, coherence analysis is highly significant over a broad range of frequencies when applied to the coupled stochastic systems. In order to be able to draw conclusion about the underlying dynamics of the cell – to –cell communication, we suggest to apply the coherence analysis to the phase fluctuations i.e. the phase of the each electrical signal subtracted by the linear trend caused by the frequency obtained by the synchronized bacterial communication system. This is motivated by the fact that phase synchronization is frequency related phenomenon. Coherence of the phase fluctuations for synchronizing oscillators is expected to be significant. For the propagation of signals the phase of both the input and the output, should be similar. This would lead to a broadband coherence between the phase fluctuations. In the case of phase synchronizing oscillators and in presence of dynamic noise, phase jumps are expected to occur. Motivated by the washboard potential of the phase difference phase jumps in both direction are expected for sufficiently large noise variance. For the signal propagation the output signal is expected to perform a phase jump after the input signal has performed one^18^.

The analysis of the bacterial ion channel can provide a fundamental insight into the structural basis of neuronal signaling. We can find that ion channels conduct long-range coherent electrical signals within the biofilm communities through spatially propagated wave of potassium and the extracellular potassium emitted from the biofilm alters the membrane potential of a distant cells. It also enables the cross species communication. This generic mechanism can attract a distinct quorum sensing bacteria towards biofilm. On the other hand, noise can play a constructive role in enhancing the synchronization of chaotic bacterial communication systems. Two uncoupled identical chaotic systems can achieve complete synchronization under the influence of a common noise. Noise can also produce amplification of a weak signal (stochastic resonance) or induce coherent oscillations in excitable and damped oscillations in membrane potential system (coherence resonance)^19^. Furthermore, noise induces a small kinetic viscosity which leads actively moving bacteria into the meta-stable states required to support quorum, given the non-local nature of the stresses mediated by autoinducers. It is to be mentioned that there are two types of noise in this kind of bacterial colony: nonlocal noise due to finite existence of bacteria and noise due to ion channel opening and closing. The role of both type of noise should be anlyzed in understanding the synchronization. This mathematical approach tends to form the patterns which is certainly based on the interaction of many components. Kinematic viscosity of the living fluid and density of quorum sensing molecules change the quorum state to non quorum state. Quorum Sensing concentration profiles, periodic and wave-like patterns^20^ can be generated out of an initially more or less non-homogeneous state. The shape of the patterns and particularly the threshold density at which the sytstem switches behaviour are therefore largely dependent on the initial condition and non-local hydrodynamics. On the otherhand, the existence of noise due to ion channel may help us to understand the long range electrical signaling through potassium waves. Essentially, the noise produced by nonlocal noise in nonlocal hydrodynamics is important in understanding the metastable state of bacteria through signaling molecules (autoinducers). This is an amazing situation where two kind of signaling are happening simultaneously: one through signaling molecules and the other through electrical signaling. It gives rise to new challenge how to formulate a comprehensive theory considering the two type of noise one in macroscopic state i.e. at the level of communication through chemical signaling and other at the level ion channel which is basically object at nanoscale.

## References

1. Sarangam Majumdar, Sukla Pal, Cross-species communication in bacterial world, J. Cell Commun. Signal. DOI 10.1007/s12079-017-0383-9 (2017).

2. Shapiro JA (1998) Thinking about bacterial populations as multicellular organisms. Annu Rev Microbiol 52: 81–104

3. Miller B. M., Bassler L. B. Quorum sensing in bacteria Annu. Rev. Microbiol., 55 (2001), pp. 165–199

4. Fuqua W. C., Winans S. C., Greenberg E. P. Quorum sensing in bacteria: the LuxR-LuxI family of cell density-responsive transcriptional regulators. J. Bacteriol. 1994;176: 269–275.

5. Majumdar, Sarangam, and Subhoshmita Mondal. “Conversation game: talking bacteria.” Journal of cell communication and signaling 10. 4 (2016): 331–335.

6. Prindle, Arthur, et al. “Ion channels enable electrical communication in bacterial communities.” Nature 527, no. 7576 (2015): 59–63.

7. Sarah D. Beagle and Steve W. Lockles., Electical signaling goes bacterial, Nature, volume 527, 2015 November.

8. Liu, Jintao, et al. “Metabolic co-dependence gives rise to collective oscillations within biofilms.” Nature 523, no. 7562 (2015): 550–554.

9. Humphries, Jacqueline, et al. “Species-Independent Attraction to Biofilms through Electrical Signaling.” Cell 168, no. 1 (2017): 200–209.

10. Lambert, Neill, et al. “Quantum biology” Nature Physics 9.1 (2013): 10–18.

11. Hille, B. Ion Channels of Excitable Membranes ( Sinauer Associates, 2001).

12. Ren, D et al. A prokaryotic voltage-gated sodium channel. Science 294, 2372–2375 (2001).

13. Iyer, R., Iverson, T. M., Accardi, A & Miller, C. A biological role for prokaryotic CIC chloride channels. Nature 419, 715–718 (2002).

14. Jiang, Y. et al. Crystal structure and mechanism of a calcium - gated potassium channel. Nature 417, 515–522 (2002).

15. Chen, G. Q., Cui, C., Mayer, M. L. & Gouaux, E. Functional characterization of a potassium–selective prokaryotic glutamate receptor. Nature 402, 817–821 (1998).

16. Pikovsky, A., Rosenblum, M. & Kurths, J. (2001) Synchronization — A Universal Concept in Nonlinear Sciences (Cambridge University Press, Cambridge).

17. Brockwell, P. J. & Davis, R. A. (1998) Time Series: Theory and Methods (Springer, NY).

18. Schelter, B. et al. Phase synchronization and coherence analysis: Sensitivity and specificity International Journal of Bifurcation and Chaos, Vol.17, No. 10 (2007) 3551–3556

19. Kiss, Z. I. et al. Noise enhanced phase synchronization and coherence resonance in sets of chaotic oscillators with weak global coupling, Chaos, 13 (1), (2003).

20. Sarangam Majumdar, Sisir Roy, Rodolfo Llinas, Bacterial “Conversations” and Pattern Formation, bioRxiv 098053, (2017).

